# Metabolic readouts of tumor instructed normal tissues (TINT) identify aggressive prostate cancer subgroups for tailored therapy

**DOI:** 10.1101/2024.04.29.591591

**Authors:** Ilona Dudka, Pernilla Wikström, Anders Bergh, Gerhard Gröbner

## Abstract

**Background:** Prostate cancer (PC) diagnosis relies on histopathological examination of prostate biopsies, which is restricted by insufficient sampling of all tumors present. Including samples from non-PC but tumor instructed normal tissues (TINT) may increase the diagnostic power by exploring the adaptive responses in benign tissues near tumors.

**Methods:** Here, we applied high-resolution magic angle spinning nuclear magnetic resonance (HR MAS NMR) to identify metabolomic biomarkers with high diagnostic value in benign prostate tissues near low/high-grade tumors.

**Results:** Benign samples near high-grade tumors (B ISUP 3+4) exhibit altered metabolic profiles compared to those close to low-grade tumors (B ISUP 1+2). The levels of six metabolites were significantly different between the two groups; myo-inositol, lysine, serine and combined signal of lysine/leucine/arginine were increased in benign samples near high-grade tumors (B ISUP 3+4) compared to near low-grade tumors (B ISUP 1+2), while levels of ethanolamine and lactate decreased. Additionally, we revealed metabolic differences in non-cancer tissues as a function of their distance to the nearest tumor. Eight metabolites (glutathione, glutamate, combined signal of glutamate/glutamine - glx, glycerol, inosine, ethanolamine, serine and arginine) significantly differentiated between benign tissue located close to the tumor (d ≤ 5 mm) compared to those far away (d ≥ 1 cm).

**Conclusions:** Our HR MAS NMR-based approach identified metabolic signatures in prostate biopsies that reflect the response of benign tissues to the presence of nearby located tumors in the same prostate and confirmed the power of the TINT concept for improved PC diagnostics and understanding of tumor-tissue interactions.

## Background

A reliable diagnosis of prostate cancer (PC) depends on a histopathological examination of biopsies (1) usually performed upon initial determination of prostate serum antigen (PSA) levels, digital rectal exam (DRE), and guided by ultrasound and magnetic resonance imaging (MRI). However, routine biopsies, due to the low prostate volume examined, sometimes miss to sample all tumors present, and they may fail to sample the most aggressive foci (2). The successful treatment of PC depends critically on the grading of all clinically significant tumor foci and the correct determination of their aggressiveness and capacity to form metastases (3). Therefore, to provide a reliable diagnostic procedure another level of potentially high information value could be included, where specific and reliable PC biomarkers in sampled non-malignant prostate tissue located near tumor foci are considered. Ideally, those nearby biomarkers should indicate not only the presence, but also the aggressiveness of tumors located elsewhere in the prostate (2).

The concept named TINT - tumor instructed normal tissue, was established by Bergh and coworkers over a decade ago (4). The TINT concept is based on a tumor induced adaptive response in histologically normal-appearing epithelium and stroma. This tumor interaction with nearby benign tissues allows subsequent growth and spreading of the tumor into the surrounding tissue environment. However, the TINT does not need to be in direct contact with the cancer epithelium and should not be mixed with the tumor stroma or the tumor microenvironment (5). TINT is also different from the so-called field cancerization or field effect, which describes pre-malignant genotypic and phenotypic alterations required for the transformation of cancer cells (6).

Recently, the diagnostic power of the TINT concept was demonstrated by monitoring the adaptation of morphologically benign prostate tissues to nearby located tumors using rat PC models (5). Additionally, in small localized PCs increased DNA synthesis was observed in remote tissue regions in a tumor type- and size-dependent manner (7). On a phenotypic level, the expression of some genes in the TINT is changed in a similar way as in tumors, while other changes seem to be exclusive to the TINT (8). Aggressive tumors may have particular effects on adjacent tissues and alterations in the non-malignant tissue located close to tumor sites can therefore provide valuable prognostic information. Low levels of phosphorylated epidermal growth factor receptor found in non-malignant and malignant prostate tissue predict favourable outcome for PC patients (2). In contrast, high levels of lysyl oxidase (LOX) in the non-malignant prostate epithelium are correlated with poor outcomes (9). Additionally, the hyaluronan staining score in the surrounding morphologically normal prostate tissue is associated with tumor aggressiveness and increased mortality risk (10). Similar correlations were found for microseminoprotein-beta (MSMB) and C/EBPβ expression levels found in surrounding tissue areas (11, 12). Moreover, in the stroma of tumors and in non-malignant prostate tissues, a high number of S100A9 positive inflammatory cells is associated with shorter cancer-specific survival (13), and a low stroma androgen receptor level is related to poor outcomes in PC patients (14).

In recent years, also metabolomic profiling has also contributed to a more comprehensive molecular understanding of PC, including understanding of the intracellular signalling pathways regulating prostate carcinogenesis (15–17). Metabolomic analysis has provided specific biomarkers for aggressive PC subtypes, which may be useful for precision medicine (18, 19). Here, we further expand the potential of the TINT concept for diagnostics and therapy by identifying metabolomic biomarkers with potentially high prognostic value in the clinic. For this purpose, we used the high resolution magic angle spinning nuclear magnetic resonance (HR MAS NMR) technique on intact prostate biopsies to provide metabolic signatures for two sets of benign prostate samples characterized by surrounding PC tumor tissue of low (B ISUP 1+2) and high (B ISUP 3+4) ISUP grades. In this way we identified six metabolites in non-cancer tissues that changed significantly as a function of the tumor grade in the surrounding tissues. Additionally, we observed that changes in eight metabolites in benign samples were related to distances to the closest tumor foci. Our findings indicate the possibility of exploiting molecular markers for reliable early diagnosis and prognosis, even in tumor-negative biopsies, which is often the case in routine clinical settings. We identified biomarkers that can differentiate between non-cancer tissues located near high-grade versus low-grade PC. Therefore, this portfolio of “TINT” biomarkers provides a powerful diagnostic tool for identifying patients with negative biopsies who are in high need of close follow-up and possibly early diagnosis and treatment.

## Materials and Methods

### Patients and tissue samples

In this study we used a well-characterized patient cohort as described previously (20). The study was conducted in accordance with the Declaration of Helsinki, and the study protocol was approved by the research ethical committee at Umeå University Hospital (Regional Ethical Review Board in Umeå). Written informed consent was obtained from each patient. Biopsy prostate samples were obtained from 31 patients treated by prostatectomy at the Urology Clinic, Umeå University Hospital, between 2009-2012. A detailed description of sample collection is described in (20). From the metabolomics profiles of 104 samples (48 benign and 56 cancer) analysed in the previous study, we selected 57 samples (27 benign and 30 cancer) from patients with unifocal cancer and additionally 35 samples (17 benign and 18 cancer) from patients with multifocal cancer, with the following conditions fulfilled: the tumor is the index tumor (highest grade and largest), the normal sample is taken from the same side, and the other tumors are located on the contralateral side and at even larger distances. The Gleason grade and ISUP grade were estimated for the cancerous tissue samples. For benign samples, Gleason grade and ISUP grade were determined based on the assignment of the closest tumor. Additionally, benign samples were characterized by distance to the closest tumor. The clinicopathological characteristics of the patients and samples are summarized in Table 1.

**Table 1.**
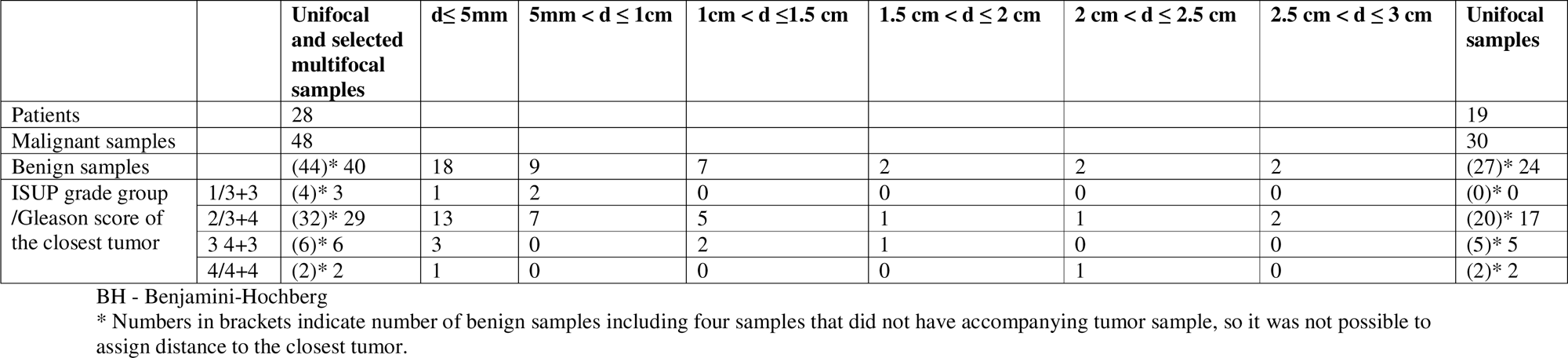
Patient and sample characteristics.

### ^1^H HR MAS MRS experiments

Metabolomic analysis of prostate tissue samples was carried out as described previously (20). Brefly, NMR experiments were carried out on a 500 MHz NMR spectrometer (Bruker Biospin, GmbH, Germany) at 277 K. We used a Carr-Purcell-Meiboom-Gill (CPMG) NMR pulse sequence with a spectral width of 20 ppm, 1024 scans, echo time of 0.2 ms, a total acquisition time of 1.64 s, a recycle delay of 1.5 s and 32 K data points. The acquired spectral data were baseline and phase corrected using TopSpin 3.6.5 software (Bruker Biospin, GmbH, Germany). Prior to peak integration seven spectra were excluded due to high levels of lipids as indicated by their pronounced NMR signals in those samples. The NMR spectra were aligned using icoshift 1.2 MATLAB 2017a (The Mathworks, Inc., USA) and the NMR peaks were integrated using an in-house developed MATLAB R2017a routine. Metabolite identification was carried out using the Chenomx NMR suite professional (version 8.6, Chenomx Inc., Edmonton, Canada).

### Multivariate analysis

Multivariate analysis of the NMR spectra was carried out in SIMCA V17 (Umetrics, Umeå, Sweden). Principal component analysis (PCA) was performed on the spectral data set to evaluate the homogeneity of the samples and identify any possible trends and outliers between the samples. Thereafter, supervised multivariate analysis orthogonal partial least squares discriminant analysis (OPLS-DA) was performed to visualize the differences between assigned groups by reducing the high dimensionality into predictive and orthogonal latent variables. Analysis of variance of cross-validated predictive residuals (CV-ANOVA) was used to assess the significance of the OPLS-DA models, where a *p* value lower than 0.05 was associated with a significant model. Leave-one-out cross validation was applied to all OPLS-DA models.

### Univariate Statistical Analysis

Metabolomic differences among assigned groups were tested by using the t-test or Mann-Whitney test for two-groups comparisons and one-way ANOVA with post hoc Benjamini-Hochberg (FDR, false discovery rate) correction for multiple comparisons (q < 0.05) for three or more group comparisons. Correlations between metabolite levels and ISUP values and distance (d) from the tumor values were calculated with Pearson correlation (two-tailed *p* value, 95% confidence interval). Univariate statistical analyses were performed with GraphPad Prism (GraphPad Software Inc., San Diego, CA, USA) version 9.4.1.

## Results

### Study description and sample characteristics

First, we aimed to identify metabolomic changes in benign prostate biopsies from PC patients associated with their location near malignant tissue of varying aggressiveness. Second, we wanted to explore variations in metabolic levels in those benign samples as a function of their distance from malignant tissue. To obtain highly reliable findings we used a well-characterized study cohort and previously obtained tissue metabolomics profiles (20). With the application of HR MAS NMR, 39 metabolites could be identified and quantified in the main NMR spectral region (0.7–8.5 ^1^H ppm). For our analysis, we included from that data set only samples from patients with unifocal cancer and from patients with multifocal cancer, where we assumed that changes in benign samples were due to the indexed tumor and not an effect of other tumor foci.

Finally, 92 samples (44 benign and 48 cancer) were used for thorough chemometric and univariate analyses. Each benign sample had additional information regarding the distance to the closest tumor and the corresponding tumor ISUP grade. For 4 samples the distance to the closest tumor was not assigned. Benign samples were divided into six groups according to the distance (d) to the tumor: group 1 (d ≤ 5 mm, 18 samples), group 2 (5 mm < d ≤ 1 cm, 9 samples), group 3 (1 cm < d ≤1.5 cm, 7 samples), group 4 (1.5 cm < d ≤ 2 cm, 2 samples), group 5 (2 cm < d ≤ 2.5 cm, 2 samples), and group 6 (2.5 cm < d ≤ 3 cm, 2 samples). In the context of ISUP grade assignment, four benign samples belonged to ISUP 1, thirty-two to ISUP 2, six to ISUP 3 and two samples to ISUP 4. Due to the limited number of samples, especially for ISUP 1 and 4, benign samples with ISUP 1 and 2 were combined into one group (B ISUP 1+2) reflecting benign samples adjacent to less-aggressive tumors, and ISUP 3 and 4 were combined into a second group (B ISUP 3+4) containing benign samples adjacent to more aggressive tumors.

Comparisons between cancer and benign samples and among different types of PC subtypes were presented in a previous paper (20). Table 1 summarizes the histological characteristics of the samples used in our final analysis.

### Benign prostate tissues classified based on ISUP grades of accompanying tumors

To understand how cancer tissue of different grades affects the metabolic fingerprint of benign samples, adequate benign samples were investigated by HR MAS NMR. The metabolic profiles of benign prostate tissue samples were modelled according to their proximity to cancer of either ISUP 1+2 or 3+4. Upon constructing corresponding PCA and OPLS-DA models with samples from patients with unifocal and multifocal cancer analysed together (R2Y = 0.224 and Q2 = 0.150, *p* = 1.97 x 10^−8^ from CV-ANOVA), the two benign groups (B ISUP 1+2 and B ISUP 3+4) could be compared with each other and with the two cancer groups (PC ISUP 1+2 and PC ISUP 3+4) as presented in Figures 1A and B. Tumor samples from the ISUP 3+4 group were generally separated from the other samples, while clear overlaps were observed between samples from other groups. However, more of the benign samples accompanying ISUP 3+4 tumors than the benign samples accompanying ISUP 1+2 tumors showed metabolic similarities with the cancer samples. When we compared all four groups using only unifocal samples, the clustering was even more clear as presented in Fig. S1 (R2Y = 0.542 and Q2 = 0.201, *p* = 0.017 from CV-ANOVA). Additionally, we observed that the two benign groups of samples showed more metabolic resemblance to their accompanying tumor groups, B ISUP 1+2 to PC ISUP 1+2 and B ISUP 3+4 to PC ISUP 3+4. Nevertheless, models comparing only two benign groups (B ISUP 1+2 *vs* B ISUP 3+4), including benign samples either from patients with unifocal and multifocal cancer analysed together or only benign samples from patients with unifocal tumors, were not significant, probably due to the limited number of benign samples with ISUP 3+4 (results not shown).

**Fig. 1.**
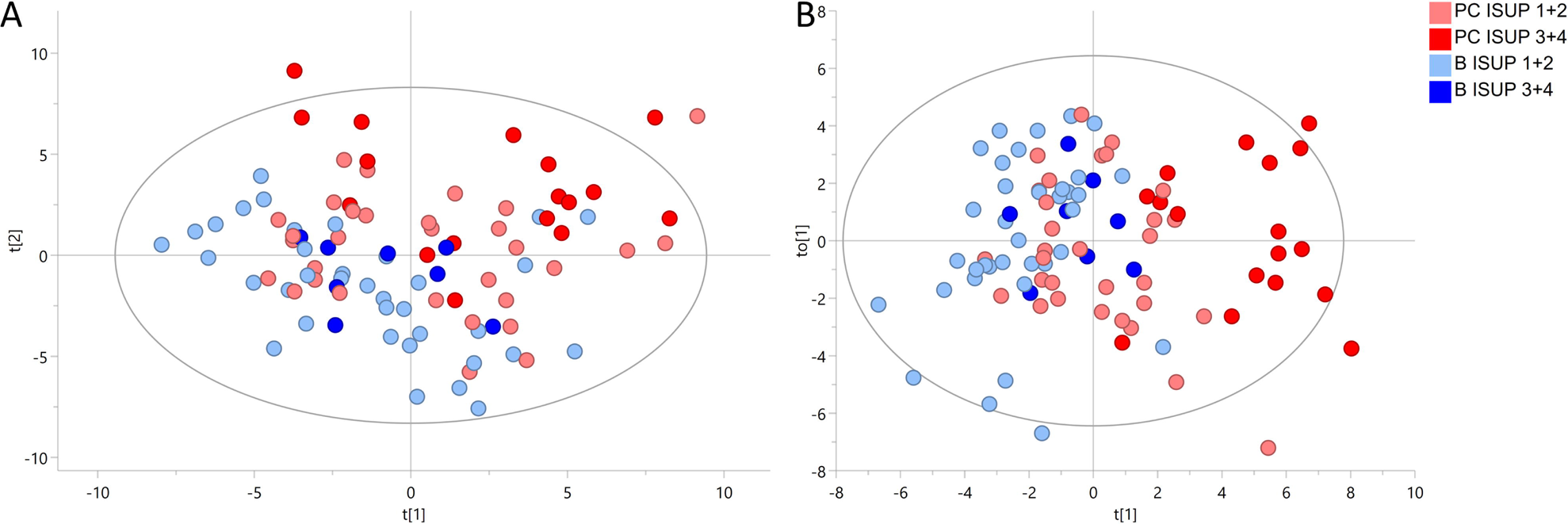
^1^H HR MAS NMR profiling analysis of samples from patients with unifocal and selected multifocal tumors among two groups of benign prostate tissues (B ISUP 1+2 and B ISUP 3+4), defined from if they were accompanied by less or more aggressive cancer foci (PC ISUP 1+2 and PC ISUP 3+4), respectively. **A** The PCA score plot. **B** The OPLS-DA score plot.

Univariate statistical analysis (t-test/Mann-Whitney test) with correction for distance from the closest tumor identified six metabolites (myo-inositol, lysine, ethanolamine, serine and lactate and combined signal of lysine/leucine/arginine) with significantly different levels between benign samples from the ISUP 1+2 and ISUP 3+4 groups (Table 2). These metabolites were also significantly correlated with the ISUP grades of the associated tumors as presented in Table 2. In addition, for those six metabolites one-way ANOVA (*p* values < 0.05) with multiple comparison tests with Benjamini-Hochberg correction was performed for four groups, two benign and two tumor (B ISUP 1+2, B ISUP 3+4, PC ISUP 1+2, PC ISUP 3+4). The results of this analysis are presented in Table 2 and in Figure 2. As shown in Fig. 2, the levels of all metabolites of benign ISUP 3+4 samples in comparison to those of the benign ISUP 1+2 group mirrored the pattern observed for tumor tissues of the ISUP 3+4 grade versus ISUP 1+2 tumors. Moreover, after plotting those metabolites separately for each benign ISUP group (Supplementary Fig. S2), increasing levels of myo-inositol, lysine and a peak for lysine/leucine/arginine starting were observed from benign samples with ISUP 1 through benign ISUP 2 and subsequent ISUP 3 grades; ultimately, ISUP 4 displayed the highest levels. The opposite trend was observed for lactate, while the levels of ethanolamine and serine did not follow a clear trend.

**Fig. 2.**
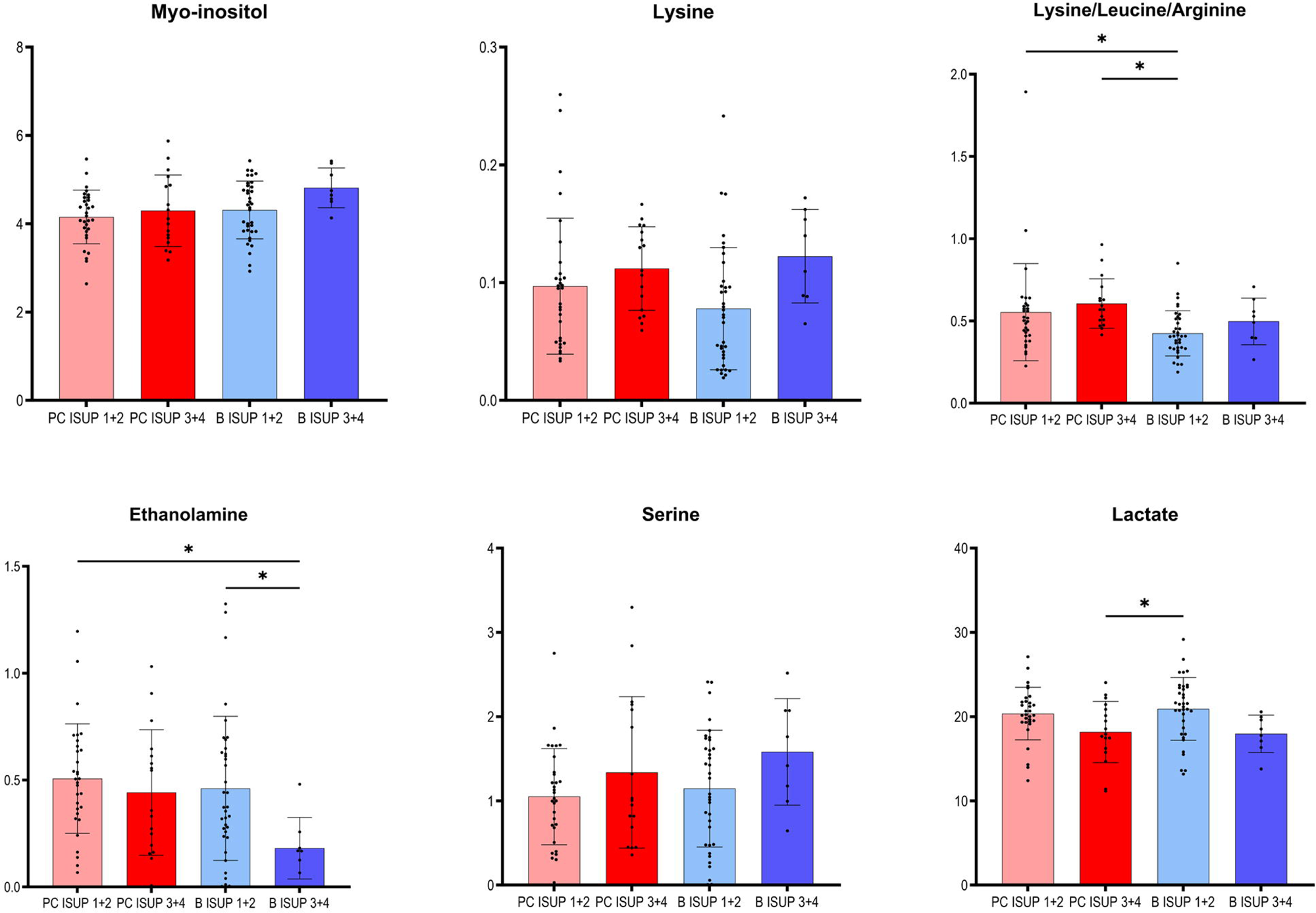
Box plots for metabolites showing significantly different levels between two groups of benign prostate tissues (B ISUP 1+2 and B ISUP 3+4), defined from if they were accompanied by less or more aggressive cancer foci (PC ISUP 1+2 and PC ISUP 3+4), respectively. Metabolites were selected based on t-test/Mann-Whitney test with correction for distance from the closest tumor. The significances marked in the figure are results of post-hoc analysis with BH correction. Analysis was based on samples from patients with unifocal and selected multifocal tumors. * *P* <[0.05.

**Table 2.** Metabolites significantly changed between benign samples accompanying tumors with ISUP 1+2 or ISUP 3+4.

| Metabolite | Correlation with ISUP values |  | B ISUP 1+2 *vs* B ISUP 3+4 * |  | One-way Anova *p*-value* PC<br>ISUP 1+2 *vs* PC ISUP 3+4 *vs*<br>B ISUP 1+2 *vs* B ISUP 3+4 | Post-hoc analysis with BH<br>correction *p*-value*<br>B ISUP 1+2 *vs* B ISUP 3+4 |
| --- | --- | --- | --- | --- | --- | --- |
|  | coefficient * | *p*-value * | *p*-value * | *p*-value after correction *<sup>c</sup> |  |  |
| Myo-inositol | 0.393/0.372 | **0.008**/0.058 | **0.047**<sup>a</sup>/0.127 | **0.037**/0.072 | 0.068 /0.135 | 0.329 /0.300 |
| Lysine | 0.428/0.365 | **0.004**/0.061 | **0.021**<sup>b</sup>/0.101<sup>a</sup> | **0.002/0.013** | **0.044** /0.110 | 0.080 /0.167 |
| Lysine/Leucine/Arginine | 0.309/0.098 | **0.041**/0.628 | 0.185<sup>a</sup>/0.800 | **0.038**/0.302 | **0.014 /0.016** | 0.487 /0.789 |
| Ethanolamine | -0.367/-0.275 | **0.014**/0.165 | **0.011**<sup>b</sup> /**0.063** | **0.036**/0.131 | 0.051 /0.250 | **0.048** /0.198 |
| Serine | 0.351/0.182 | **0.020**/0.365 | 0.110<sup>a</sup>/0.337 | **0.041**/0.239 | 0.200 /0.066 | 0.332 /0.733 |
| Lactate | -0.360/-0.259 | **0.017**/0.193 | **0.037**<sup>a</sup> /0.191 | **0.023**/0.139 | **0.018** /0.436 | 0.074 /0.533 |

BH - Benjamini-Hochberg \* First value per column was based on the analysis of benign samples from patients with unifocal and selected multifocal tumors (n = 44), the second value was based on the analysis of benign samples from patients with unifocal tumors only (n = 27). <sup>a</sup> t-test <sup>b</sup> Mann-Whitney test <sup>c</sup> correction for distance from the closest tumor

### Classification of benign prostate tissue according to the distance to the nearest tumor

To understand how distance to cancer affects the metabolic fingerprint of benign samples, adequate benign samples were investigated by HR MAS NMR. The metabolic profiles of benign prostate tissue samples were modelled according to their distance to the closest cancer foci, with tumor tissue samples also included in the model. In supplementary Figures S3 (samples from patients with unifocal and multifocal cancer analysed together) and S4 (only unifocal samples), the PCA and OPLS-DA score plots are presented for all six distance groups of benign samples together with the tumor-containing group. Here, samples from group 1 (d ≤ 5 mm) are significantly overlapped with a fraction of tumor samples, and group 3 samples (1 cm < d ≤1.5 cm) were best separated from tumor specimens. To enable a more in-depth analysis, benign samples were divided into two classification groups due to the small sample numbers, especially for groups 4 to 6. This strategy allowed a comparison of samples very close to the tumor (group 1, 0.5 cm ≤ d < 1 cm) with the combined samples from groups 2 to 6 (d ≥ 1 cm) and with tumor samples. In this way, a PCA score plot and reliable OPLS-DA model were obtained for samples from patients with both unifocal and multifocal cancer (R2Y = 0.304 and Q2Ycum = 0.161, *p* = 1.62 x 10^−4^ from CV-ANOVA) as shown in Fig. 3A and B and in Fig. S5A and B for solely unifocal samples (R2Y = 0.375 and Q2Ycum = 0.202, *p* = 1.27 x 10^−2^ from CV-ANOVA). Nevertheless, a few tumor samples still partially overlapped with the benign samples, especially those from group 1. However, models based on the two benign groups with different distances to the tumor (0.5 cm ≤ d < 1 cm vs d ≥ 1 cm) including either only unifocal or unifocal with multifocal samples, were not significant. Nevertheless, based on univariate analysis (results of t-test/Mann-Whitney test without or with correction for ISUP of accompanying tumors), key metabolites could be identified to differentiate between benign samples near a tumor (0.5 cm ≤ d < 1 cm) and samples further away (d ≥ 1 cm) as summarized in Table 3. Additionally, a Pearson correlation was performed between the assigned distance from the tumor and the levels of metabolites. One-way ANOVA with multiple comparisons test with Benjamini-Hochberg correction was used to compare the two benign groups with the third group, which included tumor samples. The results of this analysis are presented in Table 3 and Figure 4. The box plots in Fig. 4 display the obtained levels of significant metabolites found in tumors and the two groups of benign samples. As shown in Fig. S6, we also plotted the levels of those metabolites for benign samples separated into three subgroups (the first group with d = 0.5-1 cm; the second group with d = 1.5-2 cm; and the third group with d = 2.5-3 cm). Here, one can clearly explore the distance-dependent metabolic signatures of benign samples. An interesting behaviour was observed for glutathione and arginine. Glutathione levels increase with the distance from the tumor, with the lowest levels in tumor samples and the highest in benign samples 2.5 to 3.0 cm from the tumor. For arginine, the opposite trend was observed.

**Fig. 3.**
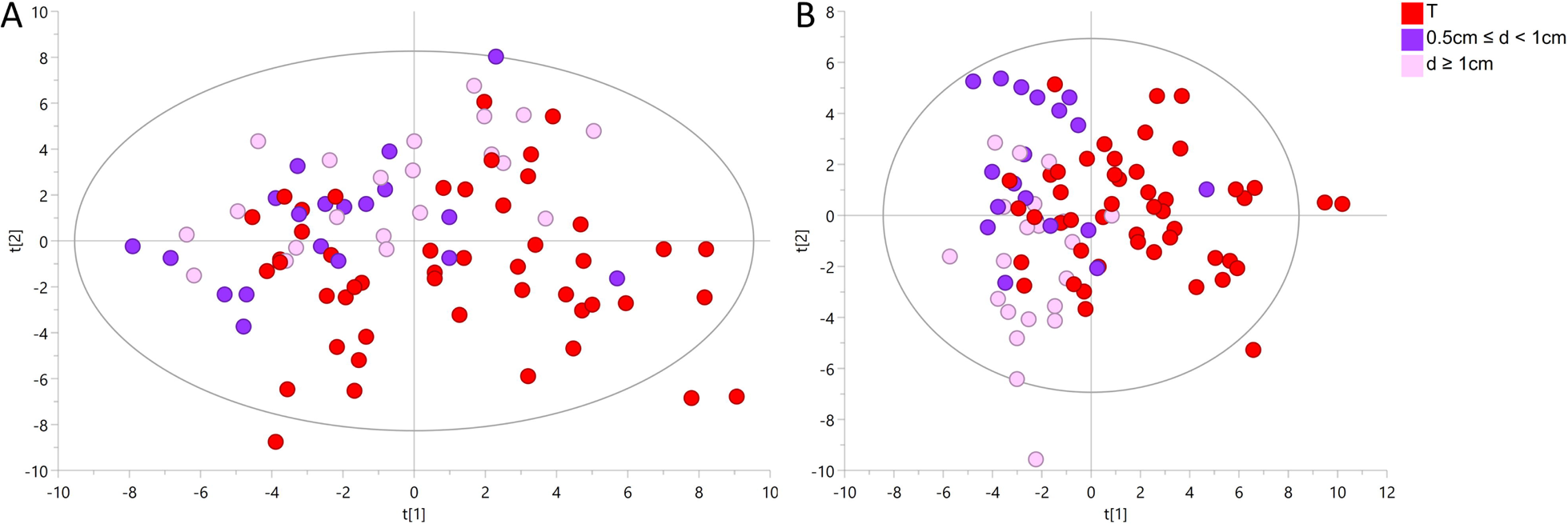
^1^H HR MAS NMR profiling analysis of two groups of benign prostate samples based on tissue distances (d) to cancer foci (group 1: 0.5 cm ≤ d < 1 cm and group 2: d ≥ 1 cm) and one group of tumor samples (T). **A** The PCA score plot of two benign groups with different distances to the closest cancer foci and one group with tumor samples. **B** The OPLS-DA score plot corresponding to the same samples as in A. Analysis was based on samples from patients with unifocal and selected multifocal tumors.

**Fig. 4.**
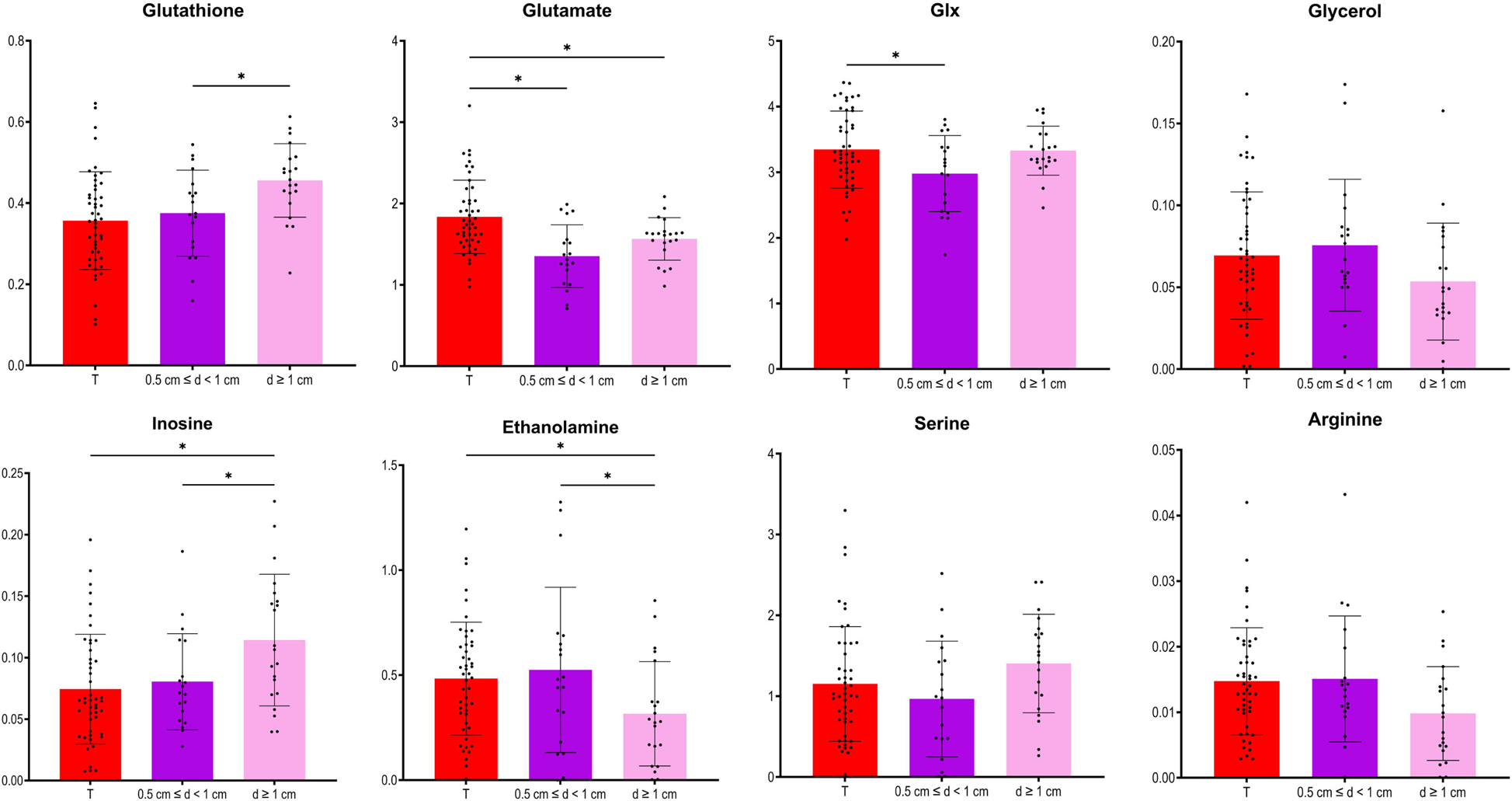
Box plots for the metabolites with significantly different levels between the two groups of benign prostate samples, close to (0.5 cm ≤ d < 1) or far away from the closest tumor (d ≥ 1 cm) presented also in tumor samples. Metabolites were selected based on t-test/Mann-Whitney test with/or without correction for distance from the closest tumor. The significances marked in the figure are results of post-hoc analysis with BH correction. Analysis was based on samples from patients with unifocal and selected multifocal tumors. * *P* <[0.05.

**Table 3.** Metabolites significantly changed with increasing distance from the closest tumor.

| Metabolite | Correlation with distance to the closest tumor |  | d ≤ 5mm vs d ≥ 1 cm |  | One -way Anova *p*-value*<br>tumor vs d ≤ 5mm vs 1 cm ≤ d ≤ 3cm | Post-hoc analysis with BH correction *p*-value*<br>d ≤ 5mm vs d ≥ 1 cm |
| --- | --- | --- | --- | --- | --- | --- |
|  | coefficient * | *p*-value* | *p*-value* | *p*-value after correction* <sup>c</sup> |  |  |
| Glutathione | 0.343/0.446 | **0.030/0.029** | **0.014<sup>a</sup> /0.022** | **0.036/0.034** | **0.004 /0.0009** | **0.037/0.032** |
| Glutamate | 0.198 /0.089 | **0.220/0.678** | **0.047<sup>a</sup> /0.470** | 0.280/0.721 | **<0,0001 /0.0003** | **0.099 /0.566** |
| Glx | 0.198/0.309 | 0.220/0.142 | **0.027<sup>a</sup> /0.071** | 0.277/0.145 | 0.041/0.130 | 0.067/0.234 |
| Inosine | 0.268/0.334 | 0.095/0.110 | **0.039<sup>b</sup> /0.013** | 0.114/0.119 | **0.005 /0.0005** | **0.033/0.010** |
| Ethanolamine | -0.349/-0.401 | **0.027/0.052** | 0.078<sup>b</sup>/0.084 | **0.036/0.053** | 0.051 /**0.037** | **0.049/0.035** |
| Serine | 0.258/0.256 | 0.109/0.228 | **0.042<sup>a</sup> /0.078** | 0.167/0.245 | 0.132/0.056 | 0.141/0.268 |
| Glycerol | -0.300/-0.391 | 0.060/0.59 | **0.031<sup>b</sup> /0.007** | 0.087/0.066 | **0.160 /0.014** | **0.176/0.021** |
| Arginine | -0.331/-0.306 | **0.037/0.146** | 0.065<sup>b</sup> /0.151 | **0.041/0.115** | 0.065/0.072 | 0.071/0.378 |

d - distance to the closest tumor
BH - Benjamini-Hochberg
\* The first value per column was based on the analysis of benign samples from patients with unifocal and selected multifocal tumors (n = 40), the second value was based on the analysis of benign samples from patients with unifocal tumors only (n = 24).
<sup>a</sup> t-test
<sup>b</sup> Mann-Whitney test
<sup>c</sup> correction for ISUP of accompanying tumors

## Discussion

Here, we identified unique metabolic signatures in prostate biopsies that reflect the response of benign tissues to the presence of nearby tumors in the same prostate. By using non-destructive HR MAS NMR spectroscopy to obtain metabolic readouts of intact biopsies, subsequent histopathological characterization can be carried out and valuable metabolic information can be obtained from histologically “normal” tissues (21). Most importantly, the obtained metabolic profiles not only presented the impact of tumors on nearby located benign tissues but also revealed that the metabolic levels of key biomarkers were sensitive to tumor grade and distance, further supporting the TINT concept (4).

Previous metabolomics studies exploring changes in benign tissues from cancerous organs have usually interpreted them as field cancerization. Yahoub et al. (21) observed significant and progressive changes in the levels of various metabolites in proximal histologically normal mucosa when comparing noncancer and cancer patients. Reed et al. (22) revealed a very strong metabolic signature differentiating normal squamous epithelium from controls and esophageal adenocarcinoma patients. Using a similar NMR approach as here, Vandergrift et al. (23) characterized the metabolomic profiles of histologically benign prostate tissues to identify tumor grade and stage and predict recurrence. In that study, elevated myo-inositol was proposed as a potential mechanistic therapeutic target in patients with highly aggressive cancer. The same group (24) also mentioned that the metabolic signatures of benign tissues of varying distances to PC lesions can differ. To identify safe margins for rectal cancer surgery, Zhang et al. (25) showed that the most significant changes in metabolite levels were observed at 0.5 cm (cT1 and cT2 stage) and 2.0 cm (cT3 and cT4 stage) from the tumor. An analogous concept was shown by Yang et al. (26) in oral squamous cell carcinoma, who identified changes in amino acids as molecular markers of the surgical margin. Jimenez et al. (27) reported that in colorectal cancer, tumor-adjacent mucosa (10 cm from the tumor margin) is characterized by unique metabolic field changes that distinguish tumors according to their T-stage and lymph node status and can accurately predict 5-year survival.

Clinically, the evaluation of PC aggressiveness is usually determined by classical histopathology of sampled prostate biopsies, including grading of observed cancer areas. The ability to detect cancer and assess aggressiveness in benign areas sampled by cancer-negative biopsies from cancer-positive patients could substantially improve the accuracy of PC diagnostics and decrease the number of unnecessary biopsies. Therefore, we focused on the concept of defining PC aggressiveness based on the metabolomic profiles of associated benign samples. The relevant OPLS-DA score plot (Fig. 1B) revealed metabolomic similarities between ISUP 1+2 PC tumors and benign samples. Notably, all significant metabolomic changes observed between benign samples from ISUP 3+4 and ISUP 1+2 groups were also noted between ISUP 3+4 and ISUP 1+2 with the levels of lactate and ethanolamine being most pronounced.

Our results clearly validate the TINT concept with the metabolomic signatures of benign prostate tissues, unambiguously reflecting the aggressiveness of nearby located tumors. The differences observed in benign samples near tumors of different aggressiveness are clearly due to cancer-driven alterations in those benign tissues. Some of the identified significant metabolites in benign samples are well established and have significant malignant potential; for example, serine showing increased levels in benign ISUP 3+4 samples compared to benign samples with ISUP 1+2. This amino acid is essential for tumor growth and progression and has been suggested as oncogenesis-supportive metabolite (28). In benign ISUP 3+4 samples, higher levels of lysine were also detected, indicating increased synthesis of collagen as described in a rat TINT model (8). Previous work indicated that the lysyl oxidase (LOX) score in TINT epithelium was correlated with tumor stage and PC survival (9). Ethanolamine was also increased in benign samples with ISUP 3+4 compared to those with ISUP 1+2. Aberrant ethanolamine phospholipid metabolism has been established as a universal metabolic hallmark of cancer (29).

In benign tissues close to aggressive tumors, the levels of myo-inositol were also elevated. This metabolite is involved in many biological processes ranging from intracellular signal transduction, calcium homeostasis regulation to energy metabolism, and it is essential for cancer formation and metastasis (35, 36). Using prostate tissue extracts, inositol hexaphosphate (IP6) was shown to significantly decrease glucose metabolism and membrane phospholipid synthesis, in addition to causing an increase in myo-inositol levels in the prostate (37). Similar changes in myo-inositol in benign prostate samples were observed by Vandergrift et al. (23). These authors suggested that myo-inositol might function as the defence response of benign tissues against nearby cancerous prostate tumors. It might thus provide endogenous tumor suppression of aggressive PC growth. Overall, elevations in the osmolyte, myo-inositol, may indicate localized changes in osmoregulation (38).

In this study, we also found metabolic changes as a function of the distance of benign samples to the next tumor locus. Here, eight metabolites were significantly different between benign tissue samples in collected close to the tumor (d ≤ 5 mm) compared to those far away from the tumor (d ≥ 1 cm). By further clustering benign samples every 5 mm from the tumor, three differently behaving benign groups could be identified. Moreover, four of those metabolites showed the largest differences between tumor and benign samples in the d = 1.5-2 cm distance region. Interestingly, level of glutathione in the benign samples gradually increased as the distance from the tumor increased. Glutathione is the primary cellular antioxidant and plays a key role in carcinogenesis and the modulation of the cellular response to antineoplastic agents (39). Therefore, molecular changes in the glutathione antioxidant system and disturbances in glutathione homeostasis are implicated in tumor initiation, progression, and treatment response. Although removal and detoxification of carcinogens are crucial in healthy cells, elevated glutathione levels in tumor cells are associated with tumor progression and increased resistance to chemotherapeutic drugs (39). Our observation that glutathione levels increased with distance is consistent with our previous reports of downregulation of glutathione S-transferases in rat TINT models (8).

Dysregulation of the oxidative pathway was also visible by decreased levels of arginine and increased levels of combined signal of glutamate/glutamine (glx) and glutamate (a precursor of glutathione) in benign samples more remote from tumor locations. The observed deviation in glutamine metabolism is typical for cancer cells and drives their growth and cancer progression. PC is a tumor type that is heavily dependent on glutamine for growth and survival (30, 31). Interestingly, arginine showed an opposite trend relative to glutathione, with the highest levels in tumor samples and decreasing levels in benign samples as a function of their distance to the tumor. Arginine deprivation has been used to target cancers (32), and its metabolism affects not only malignant cells but also surrounding immune cells (33).

## Conclusion

In summary, HR MAS NMR-based metabolomics of intact biopsies is shown here as a powerful approach for identifying metabolomic alterations in benign biopsies from cancerous prostates. This approach not only provides additional valuable information not accessible by standard clinical histopathological assignment but can also be used for improved diagnostics of PC. In addition, the outcome of our study provides clear evidence for the TINT concept that tumor indicating normal tissues are also affected at the metabolomic level. Moreover, the degree of metabolomic alteration reflects even the aggressiveness of neighbouring tumor and the distance from it. Therefore, the HR MAS NMR analysis approach of prostate biopsies has important clinical potential for identifying cancer and its aggressive state in benign biopsies from cancerous prostates. Understanding the dynamics and mechanisms driving the cross-talk between tumor and non-cancerous “benign” tissue in the affected prostate aid in understanding prostate carcinogenesis, progression, recurrence, and resistance to treatment. This can provide new opportunities for the diagnosis and prognostication of PC patients.

## Abbreviations

CPMG: Carr-Purcell-Meiboom-Gill
DRE: digital rectal exam
Glx: glutamate/glutamine
HR MAS NMR: High Resolution Magic Angle Spinning Nuclear Magnetic Resonance
IP6: inositol hexaphosphate
ISUP: International Society of Urological Pathology
LOX: lysyl oxidase
MSMB: microseminoprotein-beta
PC: Prostate cancer
PCA: Principal Component Analysis
PSA: Prostate-Specific Antigen
TINT: tumor instructed normal tissues

## Ethics approval and consent to participate

This study was conducted in accordance with the Declaration of Helsinki, and the study protocol was approved by the research ethical committee at Umeå University Hospital (Regional Ethical Review Board in Umeå). Written informed consent was obtained from each patient.

## Consent for publication

All authors have reviewed the final version of the manuscript and approved it for publication.

## Availability of data and materials

The datasets used or analysed during the current study are available from the corresponding authors on reasonable request.

## Competing interests

The authors declare that no competing interests exist.

## Funding

This work was supported by grants from Swedish Research Council (2022-00946, 2021-06146), the Swedish Cancer Society (21-1856 and 22-2041), the Kempe Foundation, the Knut and Alice Wallenberg Foundation (“NMR for Life” Programme), the SciLifeLab, SwedNMR with the Swedish National NMR Centre and Umeå Insamlingsstiftelse. The funding bodies were not involved in the design of this study, in the collection, analysis, and interpretation of the data, or in the writing of the manuscript.

## Author contributions

Conception and design: PW, AB, GG; Acquisition of data: ID, AB, GG; Analysis and interpretation of data: ID, KL, PW, AB, GG; Writing, review, and/or revision of the manuscript: ID, PW, AB, GG; Study supervision: PW, AB, GG. All the authors have read and approved the final manuscript.

## Acknowledgements

Skillful technical assistance was provided by Mrs. Pernilla Andersson and Susanne Gidlund.

## Supplementary material

**Additional file 1: Fig. S1** ^1^H HR MAS NMR profiling analysis of samples from patients with only unifocal tumors among two groups of benign prostate tissues (B ISUP 1+2 and B ISUP 3+4), defined from if they were accompanying by less or more aggressive cancer foci (PC ISUP 1+2 and PC ISUP 3+4). **A** The PCA score plot. **B** The OPLS-DA score plot.

**Additional file 2: Fig. S2** Box plots for the selected metabolites significantly different between the two groups of benign prostate tissues accompanying less-aggressive tumors (B ISUP 1+2) and benign prostate tissues accompanying more-aggressive tumors (B ISUP 3+4) presented for four benign ISUP groups, B ISUP 1, B ISUP 2, B ISUP 3, B ISUP 4. Metabolites were selected based on t-test/Mann-Whitney test with correction for distance from the closest tumor. Analysis was based on samples from patients with unifocal and selected multifocal tumors.

**Additional file 3: Fig. S3** ^1^H HR MAS NMR profiling analysis of samples from patients with unifocal and selected multifocal tumors among tumor samples and six groups of benign prostate samples depending on the distance to the closest tumor, where group 1 (0.5 cm ≤ d < 1 cm), group 2 (d = 1 cm), group 3 (d = 1.5 cm), group 4 (d = 2.0 cm), group 5 (d = 2.5 cm), and group 6 (d = 3.0 cm). **A** The PCA score plot. **B** The OPLS-DA score plot.

**Additional file 4: Fig. S4** ^1^H HR MAS NMR profiling analysis of only samples from patients with unifocal tumor among tumor samples and six groups of benign prostate samples depending on the distance to the closest tumor, where group 1 (0.5 cm ≤ d < 1 cm), group 2 (d = 1 cm), group 3 (d = 1.5 cm), group 4 (d = 2.0 cm), group 5 (d = 2.5 cm), and group 6 (d = 3.0 cm). **A** The PCA score plot. **B** The OPLS-DA score plot.

**Additional file 5: Fig. S5** ^1^H HR MAS NMR profiling analysis of two groups of benign prostate samples based on tissue distances (d) to cancer foci (group 1: 0.5 cm ≤ d < 1 cm and group 2: d ≥ 1 cm) and one group of tumor samples. **A** The PCA score plot of two benign groups with different distances to the closest cancer foci and one group with tumor samples. **B** The OPLS-DA score plot corresponding to the same samples as in A. Analysis was based on samples from patients with only unifocal tumors.

**Additional file 6: Fig. S6** Box plots for the selected metabolites significantly different between the two groups of benign prostate tissues, close to (0.5 cm ≤ d < 1) or far away from the closest tumor (d ≥ 1 cm) presented for three benign distance groups, group 1 (0.5cm ≤ d ≤ 1 cm), group 2 (1.5 cm ≤ d ≤ 2 cm), group 3 (2.5 cm ≤ d ≤ 3 cm) and group of tumor samples. Metabolites were selected based on t-test/Mann-Whitney test with/or without correction for distance from the closest tumor. The significances marked in the figure are results of post-hoc analysis with BH correction. Analysis was based on samples from patients with unifocal and selected multifocal tumors. * P < 0.05.

## Notes

### Competing Interest Statement

The authors have declared no competing interest.

## References

1. Mottet N, van den Bergh RCN, Briers E, Van den Broeck T, Cumberbatch MG, De Santis M, et al. EAU-EANM-ESTRO-ESUR-SIOG Guidelines on Prostate Cancer-2020 Update. Part 1: Screening, Diagnosis, and Local Treatment with Curative Intent. Eur Urol. 2021;79(2):243–62.

2. Hammarsten P, Karalija A, Josefsson A, Rudolfsson SH, Wikstrom P, Egevad L, et al. Low levels of phosphorylated epidermal growth factor receptor in nonmalignant and malignant prostate tissue predict favorable outcome in prostate cancer patients. Clin Cancer Res. 2010;16(4):1245–55.

3. Teo MY, Rathkopf DE, Kantoff P. Treatment of Advanced Prostate Cancer. Annu Rev Med. 2019;70:479–99.

4. Halin S, Hammarsten P, Adamo H, Wikström P, Bergh A. Tumor indicating normal tissue could be a new source of diagnostic and prognostic markers for prostate cancer. Expert Opinion on Medical Diagnostics. 2010;5(1):37–47.

5. Adamo HH, Stromvall K, Nilsson M, Bergstrom SH, Bergh A. Adaptive (TINT) Changes in the Tumor Bearing Organ Are Related to Prostate Tumor Size and Aggressiveness. Plos One. 2015;10(11).

6. Curtius K, Wright NA, Graham TA. An evolutionary perspective on field cancerization. Nat Rev Cancer. 2018;18(1):19–32.

7 . Halin Bergstrom S, Lundholm M, Nordstrand A, Bergh A. Rat prostate tumors induce DNA synthesis in remote organs. Sci Rep. 2022;12(1):7908.

8. Adamo HH, Bergstrom SH, Bergh A. Characterization of a Gene Expression Signature in Normal Rat Prostate Tissue Induced by the Presence of a Tumor Elsewhere in the Organ. Plos One. 2015;10(6).

9. Nilsson M, Hagglof C, Hammarsten P, Thysell E, Stattin P, Egevad L, et al. High Lysyl Oxidase (LOX) in the Non-Malignant Prostate Epithelium Predicts a Poor Outcome in Prostate Cancer Patient Managed by Watchful Waiting. Plos One. 2015;10(10).

10. Josefsson A, Adamo H, Hammarsten P, Granfors T, Stattin P, Egevad L, et al. Prostate Cancer Increases Hyaluronan in Surrounding Nonmalignant Stroma, and This Response Is Associated with Tumor Growth and an Unfavorable Outcome. Am J Pathol. 2011;179(4):1961–8.

11. Bergstrom SH, Jaremo H, Nilsson M, Adamo HH, Bergh A. Prostate tumors downregulate microseminoprotein-beta (MSMB) in the surrounding benign prostate epithelium and this response is associated with tumor aggressiveness. Prostate. 2018;78(4):257–65.

12. Adamo H, Hammarsten P, Hägglöf C, Dahl Scherdin T, Egevad L, Stattin P, et al. Prostate cancer induces C/EBPβ expression in surrounding epithelial cells which relates to tumor aggressiveness and patient outcome. The Prostate. 2018;79(5):435–45.

13. Tidehag V, Hammarsten P, Egevad L, Grantors T, Stattin P, Leanderson T, et al. High density of S100A9 positive inflammatory cells in prostate cancer stroma is associated with poor outcome. Eur J Cancer. 2014;50(10):1829–35.

14. Wikstrom P, Marusic J, Stattin P, Bergh A. Low Stroma Androgen Receptor Level in Normal and Tumor Prostate Tissue Is Related to Poor Outcome in Prostate Cancer Patients. Prostate. 2009;69(8):799–809.

15. Rye MB, Krossa S, Hall M, van Mourik C, Bathen TF, Drablos F, et al. The genes controlling normal function of citrate and spermine secretion are lost in aggressive prostate cancer and prostate model systems. iScience. 2022;25(6):104451.

16. Andersen MK, Hoiem TS, Claes BSR, Balluff B, Martin-Lorenzo M, Richardsen E, et al. Spatial differentiation of metabolism in prostate cancer tissue by MALDI-TOF MSI. Cancer Metab. 2021;9(1):9.

17. Bruzzone C, Loizaga-Iriarte A, Sanchez-Mosquera P, Gil-Redondo R, Astobiza I, Diercks T, et al. (1)H NMR-Based Urine Metabolomics Reveals Signs of Enhanced Carbon and Nitrogen Recycling in Prostate Cancer. J Proteome Res. 2020;19(6):2419–28.

18. Hansen AF, Hoiem TS, Selnaes KM, Bofin AM, Storkersen O, Bertilsson H, et al. Prediction of recurrence from metabolites and expression of TOP2A and EZH2 in prostate cancer patients treated with radiotherapy. Nmr Biomed. 2023;36(5):e4694.

19. Dudka I, Thysell E, Lundquist K, Antti H, Iglesias-Gato D, Flores-Morales A, et al. Comprehensive metabolomics analysis of prostate cancer tissue in relation to tumor aggressiveness and TMPRSS2-ERG fusion status. BMC Cancer. 2020;20(1):437.

20. Dudka I, Lundquist K, Wikstrom P, Bergh A, Grobner G. Metabolomic profiles of intact tissues reflect clinically relevant prostate cancer subtypes. J Transl Med. 2023;21(1):860.

21. Yakoub D, Keun HC, Goldin R, Hanna GB. Metabolic Profiling Detects Field Effects in Nondysplastic Tissue from Esophageal Cancer Patients. Cancer Res. 2010;70(22):9129–36.

22. Reed MAC, Singhal R, Ludwig C, Carrigan JB, Ward DG, Taniere P, et al. Metabolomic Evidence for a Field Effect in Histologically Normal and Metaplastic Tissues in Patients with Esophageal Adenocarcinoma. Neoplasia. 2017;19(3):165–74.

23. Vandergrift LA, Decelle EA, Kurth J, Wu S, Fuss TL, DeFeo EM, et al. Metabolomic Prediction of Human Prostate Cancer Aggressiveness: Magnetic Resonance Spectroscopy of Histologically Benign Tissue. Sci Rep. 2018;8(1):4997.

24. Dinges SS, Vandergrift LA, Wu S, Berker Y, Habbel P, Taupitz M, et al. Metabolomic prostate cancer fields in HRMAS MRS-profiled histologically benign tissue vary with cancer status and distance from cancer. Nmr Biomed. 2019;32(10):e4038.

25. Zhang S, Pan G, Liu Z, Kong Y, Wang D. Association of levels of metabolites with the safe margin of rectal cancer surgery: a metabolomics study. BMC Cancer. 2022;22(1):1043.

26. Yang XH, Zhang XX, Jing Y, Ding L, Fu Y, Wang S, et al. Amino acids signatures of distance-related surgical margins of oral squamous cell carcinoma. EBioMedicine. 2019;48:81–91.

27. Jimenez B, Mirnezami R, Kinross J, Cloarec O, Keun HC, Holmes E, et al. 1H HR-MAS NMR spectroscopy of tumor-induced local metabolic “field-effects” enables colorectal cancer staging and prognostication. J Proteome Res. 2013;12(2):959–68.

28. Geeraerts SL, Heylen E, De Keersmaecker K, Kampen KR. The ins and outs of serine and glycine metabolism in cancer. Nat Metab. 2021;3(2):131–41.

29. Cheng M, Bhujwalla ZM, Glunde K. Targeting Phospholipid Metabolism in Cancer. Front Oncol. 2016;6:266.

30. Ni R, Li ZW, Li L, Peng D, Ming Y, Li L, et al. Rethinking glutamine metabolism and the regulation of glutamine addiction by oncogenes in cancer. Frontiers in Oncology. 2023;13.

31. Lasorsa F, di Meo NA, Rutigliano M, Ferro M, Terracciano D, Tataru OS, et al. Emerging Hallmarks of Metabolic Reprogramming in Prostate Cancer. Int J Mol Sci. 2023;24(2).

32. Chen CL, Hsu SC, Chung TY, Chu CY, Wang HJ, Hsiao PW, et al. Arginine is an epigenetic regulator targeting TEAD4 to modulate OXPHOS in prostate cancer cells. Nat Commun. 2021;12(1).

33. Matos A, Carvalho M, Bicho M, Ribeiro R. Arginine and Arginases Modulate Metabolism, Tumor Microenvironment and Prostate Cancer Progression. Nutrients. 2021;13(12).

34. Lounis MA, Ouellet V, Peant B, Caron C, Li ZH, Al-Mass A, et al. Elevated Expression of Glycerol-3-Phosphate Phosphatase as a Biomarker of Poor Prognosis and Aggressive Prostate Cancer. Cancers. 2021;13(6).

35. Chen Q, Shen LF, Li S. Emerging role of inositol monophosphatase in cancer. Biomed Pharmacother. 2023;161.

36. Case KC, Schmidtke MW, Greenberg ML. The paradoxical role of inositol in cancer: a consequence of the metabolic state of a tumor. Cancer Metast Rev. 2022;41(2):249–54.

37. Raina K, Ravichandran K, Rajamanickam S, Huber KM, Serkova NJ, Agarwal R. Inositol Hexaphosphate Inhibits Tumor Growth, Vascularity, and Metabolism in TRAMP Mice: A Multiparametric Magnetic Resonance Study. Cancer Prev Res. 2013;6(1):40–50.

38. Tian Y, Nie X, Xu S, Li Y, Huang T, Tang HR, et al. Integrative metabonomics as potential method for diagnosis of thyroid malignancy. Sci Rep-Uk. 2015;5.

39. Kennedy L, Sandhu JK, Harper ME, Cuperlovic-Culf M. Role of Glutathione in Cancer: From Mechanisms to Therapies. Biomolecules. 2020;10(10).

40. Kim IS, Jo EK. Inosine: A bioactive metabolite with multimodal actions in human diseases. Front Pharmacol. 2022;13.

