## Supplementary figures and images for "Metabolic readouts of tumor instructed normal tissues (TINT) identify aggressive prostate cancer subgroups for tailored therapy"

### Supplementary Figure S1

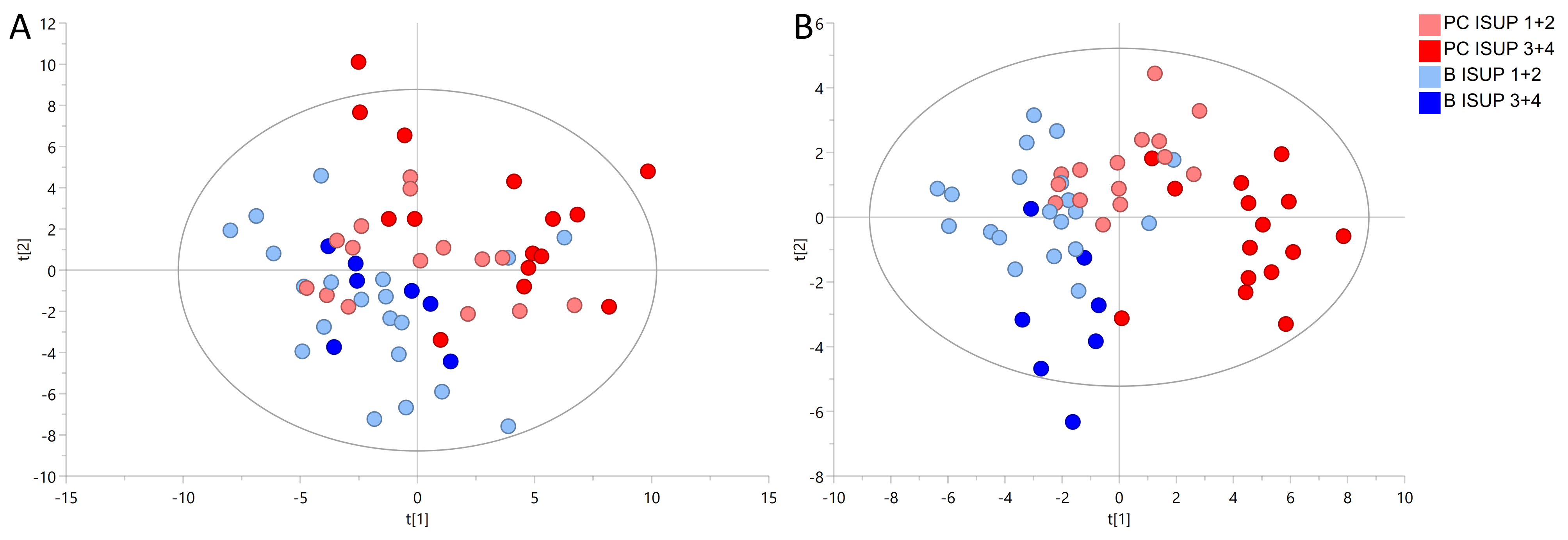

### Supplementary Figure S2

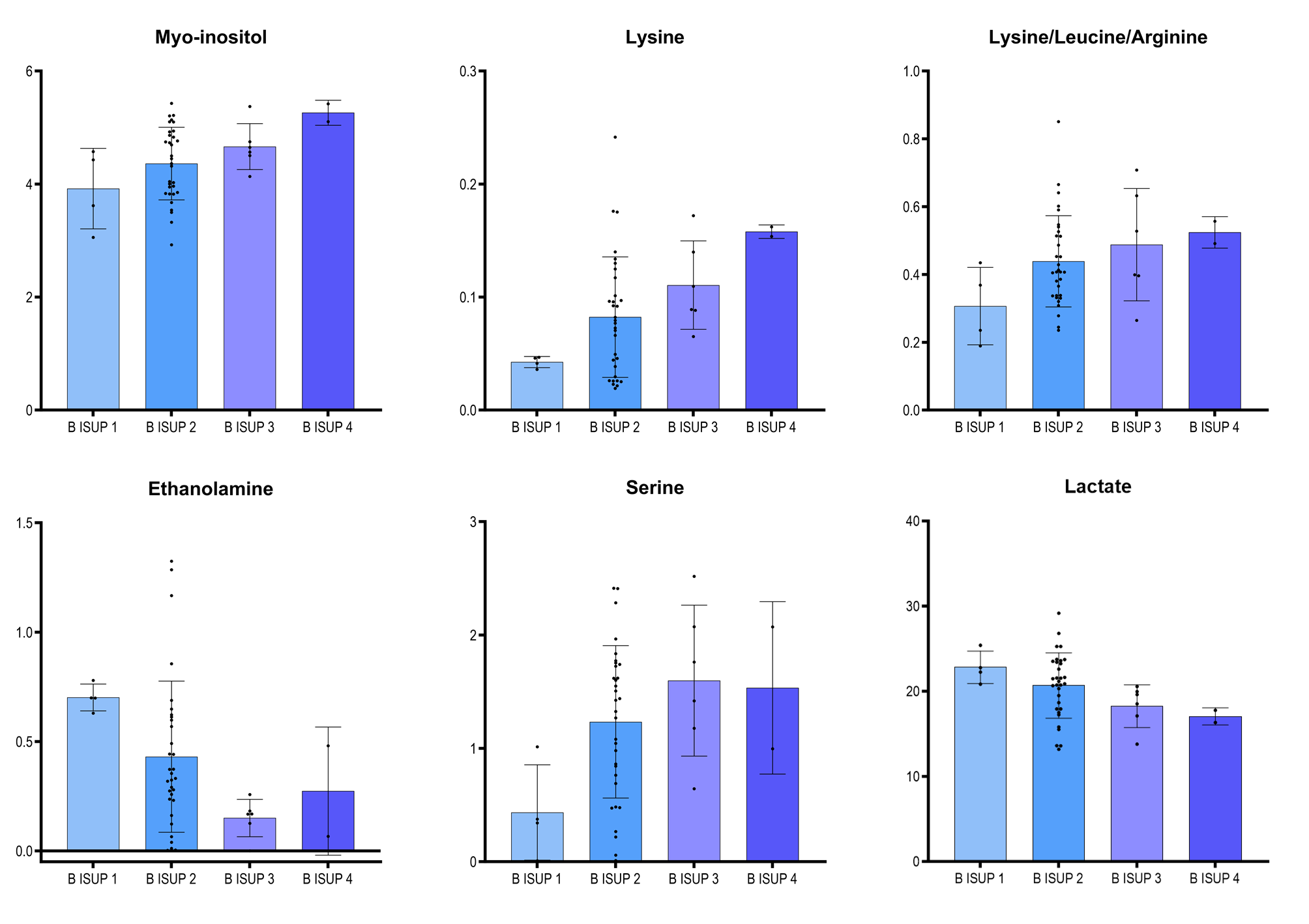

### Supplementary Figure S3

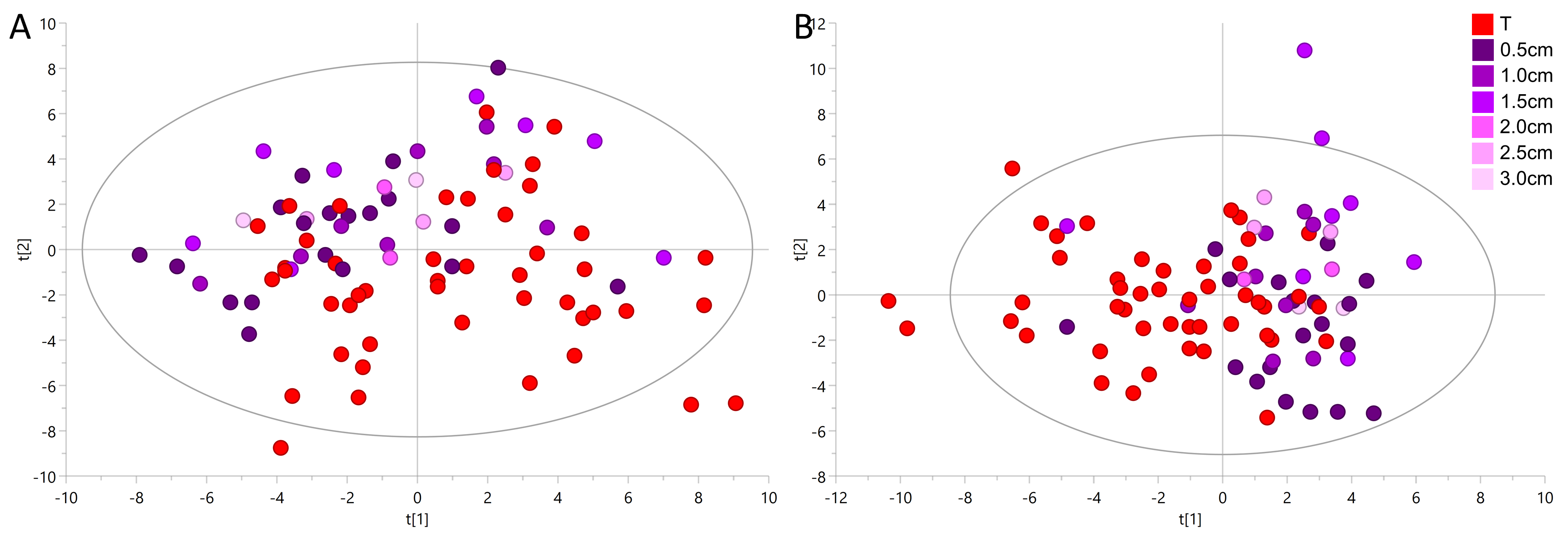

### Supplementary Figure S4

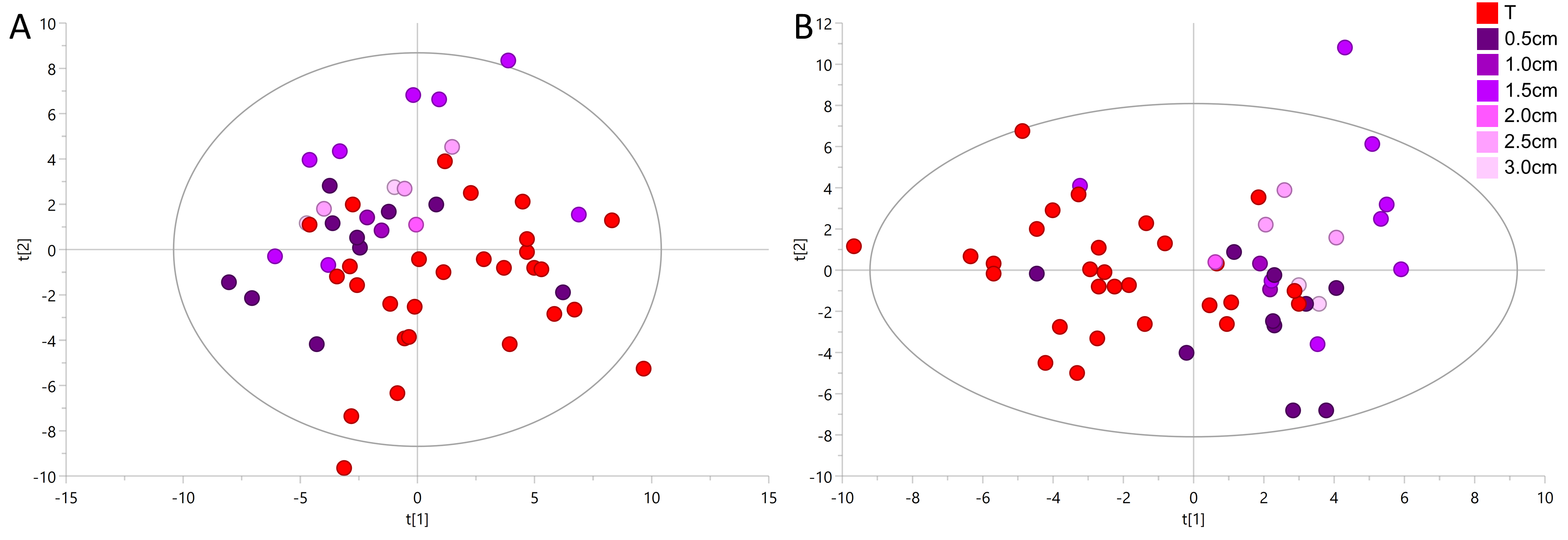

### Supplementary Figure S5

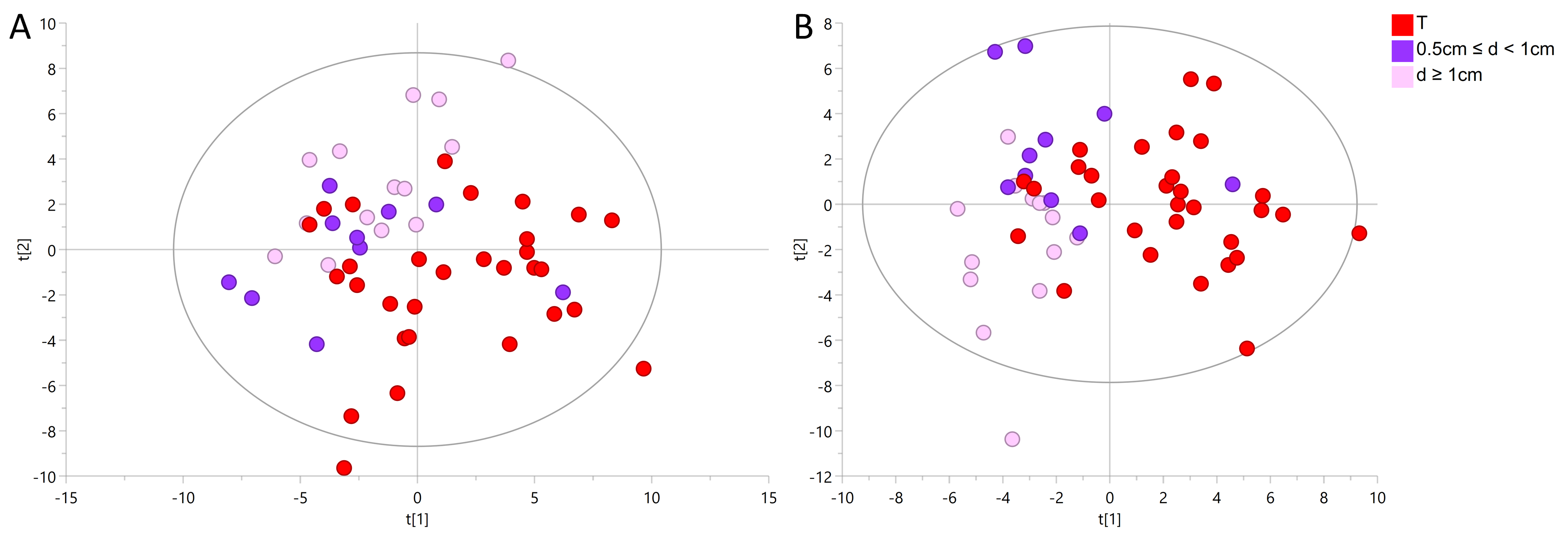

### Supplementary Figure S6

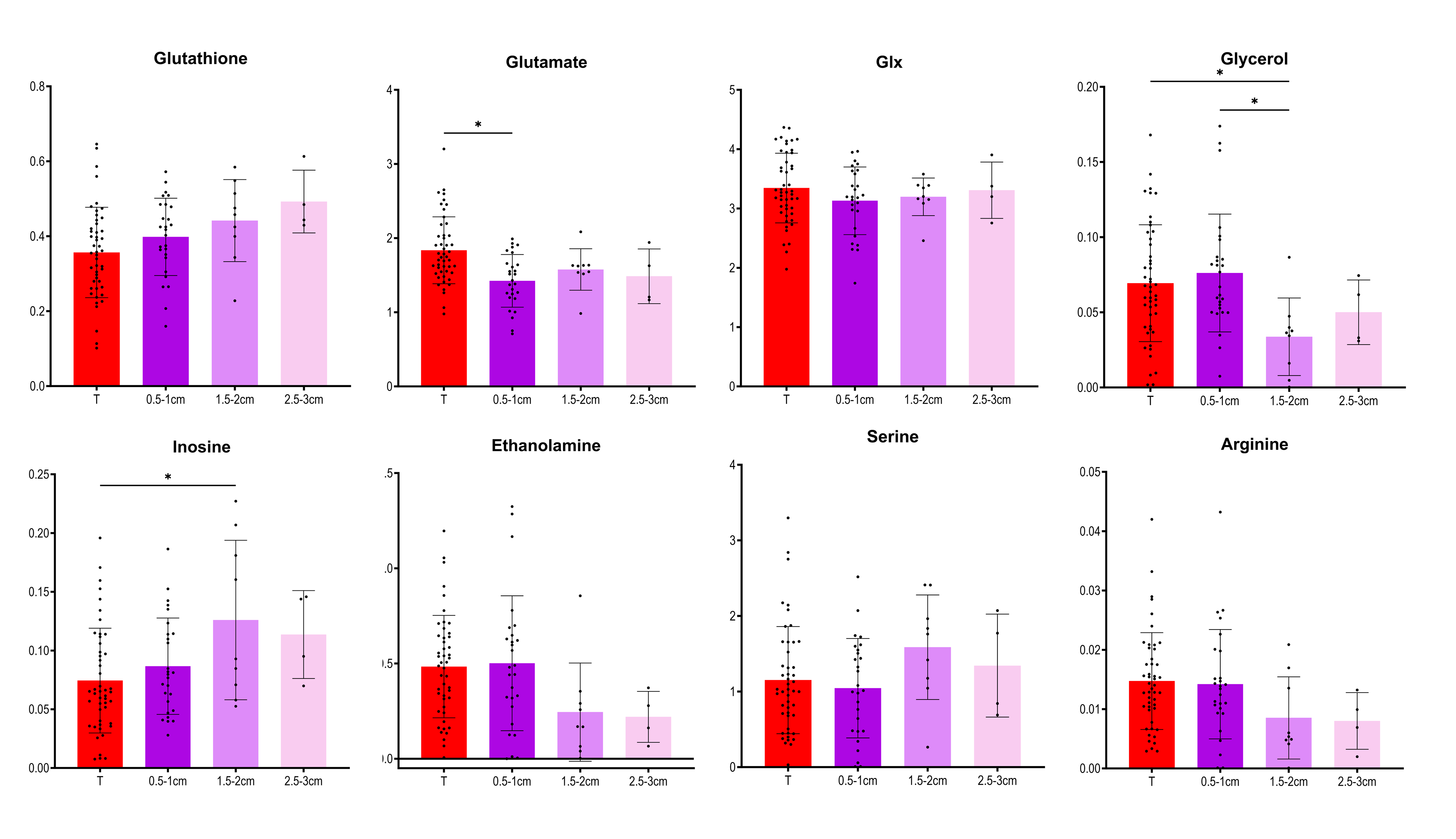
